# Cones and Cone Pathways Remain Functional in Advanced Retinal Degeneration

**DOI:** 10.1101/2022.09.27.509752

**Authors:** Erika M. Ellis, Antonio E. Paniagua, Miranda L. Scalabrino, Mishek Thapa, Jay Rathinavelu, Yuekan Jiao, David S. Williams, Greg D. Field, Gordon L. Fain, Alapakkam P. Sampath

**Author notes:** Correspondence (A.P.S.), (G.L.F.), (G.D.F.), (D.S.W.).

## Abstract

Most defects causing retinal degeneration in retinitis pigmentosa (RP) are rod-specific mutations, but the subsequent degeneration of cones, which produces loss of daylight vision and high-acuity perception, is the most debilitating feature of the disease. To understand better why cones degenerate and how cone vision might be restored, we have made the first single-cell recordings of light responses from degenerating cones and retinal interneurons after most rods have died and cones have lost their outer-segment disk membranes and synaptic pedicles. We show that degenerating cones have functional cyclic-nucleotide-gated channels and can continue to give light responses, apparently produced by opsin localized either to small areas of organized membrane near the ciliary axoneme or distributed throughout the inner segment. Light responses of second-order horizontal and bipolar cells are smaller and less sensitive but surprisingly similar to those of normal retina. Furthermore, retinal output as reflected in responses of ganglion cells is largely intact at cone-mediated light levels. Together, these findings show that cones and their retinal pathways can remain functional even as degeneration is progressing, an encouraging result for future research aimed at enhancing the light sensitivity of residual cones to restore vision in patients with genetically inherited retinal degeneration.

## INTRODUCTION

Retinitis pigmentosa (RP) is a group of inherited retinal degenerative diseases producing progressive photoreceptor death, vision loss, and eventual blindness [1]. The majority of genes associated with RP are rod-specific genes, but it is the secondary death of cones that leads to loss of high-acuity vision and blindness. Cone death has been shown to begin at about the time rod degeneration is mostly complete [2]. Cones first lose their outer segments, then axon and synaptic terminal (pedicle), and finally most of the inner segment to form rounded cell bodies [3-5]. These cones, sometimes called “dormant”, can remain viable for a prolonged period of at least a year in mouse [6] but several years in pig [7], often sprouting telodendria throughout the remnant of the outer nuclear layer [5].

Although many possible explanations have been given for why cones die after the death of the rods, recent work has focused on a lack of metabolic energy [2, 7-11]. Since the most energetically demanding process in a photoreceptor is the maintenance of ion gradients required to generate membrane currents [12, 13], we wondered whether metabolic dysfunction would alter ion channels leading to the loss of light responses and voltage-gated Ca^2+^ currents responsible for synaptic transmission. At present, little is known about changes in the cellular and synaptic physiology of the retina during photoreceptor degeneration.

We have therefore made patch-clamp and multi-electrode-array recordings from retinal cells in the central region of the retina of rd10 mice, after rods have mostly died and cones have lost their outer segments and synaptic pedicles. We have made the surprising discovery that cones continue to respond to light, and that light responses of second-order horizontal and bipolar cells are smaller and less sensitive but otherwise surprisingly similar to those in normal retina. The signal fidelity of retinal ganglion cells (RGCs) is substantially diminished under scotopic and mesopic conditions but very similar to that in healthy retinas in brighter light, with only a moderate decrease in photopic responsiveness. Furthermore, spatiotemporal receptive fields at cone-mediated light levels remain comparable in structure to those of healthy retinas. We conclude that degenerating cones do not shut down to preserve energy and are not dormant but, on the contrary, continue with the rest of the retina to respond to light and remain functional for as long as possible [14]. These results give support to efforts to reactivate cones by enhancing their light sensitivity, in the hope of restoring nearly normal photopic vision in patients with degenerative disease.

## RESULTS

To examine the function of cone-mediated vision in relatively late-stage RP, we targeted experiments to ages at which rods were nearly all dead and nearly all cones had undergone extensive morphological changes in rd10 mice. Previous work has shown that by 4 weeks of age, rd10 mice have many fewer rods and cones with a deformed shape [15-18]. By 6 weeks postnatal, nearly all the rods have degenerated, and the outer segments and synaptic pedicles of the cones were no longer observable in central retina where we made our recordings, leaving cones with only a small inner segment and nucleus (Figure 1A). At 9 weeks, nearly all the rods and many cones were gone, but in surviving cones we could often detect processes emerging from the proximal end of the inner segment, resembling axonal telodendria or synaptic dendrites as others have previously observed [for example 5].

**Figure 1.**
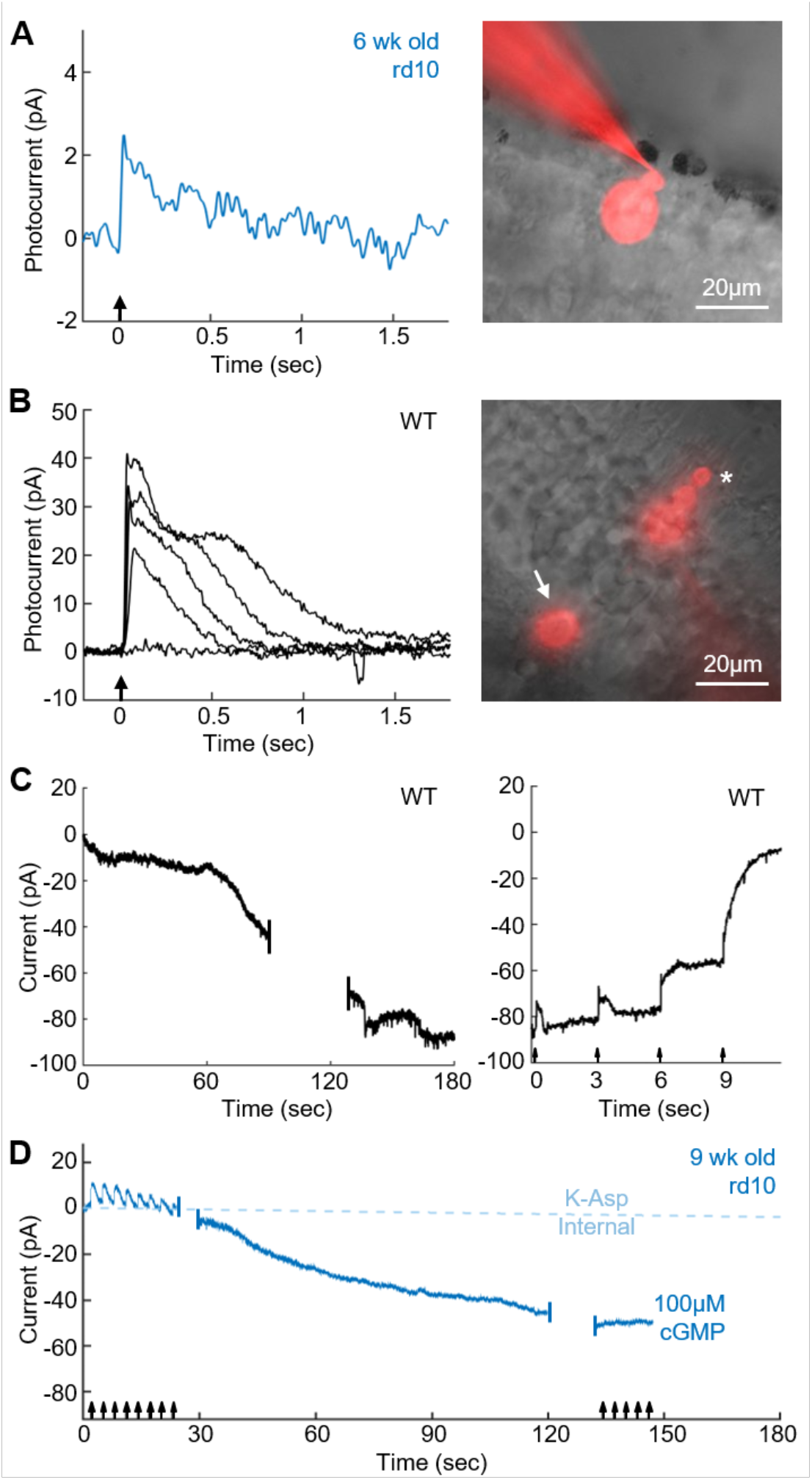
Degenerating cones maintain expression of CNG channels. Light responses (left) and images (right) from a degenerating cone in 6-week-old rd10 retina (A) and WT cone (B); note difference of scale for photocurrents. Arrows indicate 10ms flashes from 405-nm LED delivering 2.7×10^7^ photons μm^−2^ (A); and 3.0×10^2^, 4.0×10^3^, 1.5 ×10^4^, 4.0×10^4^, and 1.5×10^5^ photons μm^−2^ (B). Images of cells filled with Alexa-750 are oriented with the photoreceptor layer at the top right. In A, note that outer nuclear layer was almost entirely degenerated with only a few cone cell bodies remaining. Outer segment and cone pedicle absent in A but clearly visible in the WT cone (asterisk and arrow in B). Scale bars, 20μm. (C) Change in current (V_m_ = −40mV) with whole-cell dialysis of 100 μM cGMP into WT cone. Left, initial development of current as cGMP diffused into cell. Right at steady state, 10 ms flashes from a 405-nm LED (arrows) of successively increasing intensities of 6.0×10^2^, 7.5×10^3^, 8.0×10^4^, and 2.5×10^5^ photons μm^−2^. (D) cGMP dialysis in 9-week-old rd10 retina. Cone was periodically stimulated with flashes delivering 2.7×10^7^ photons μm^−2^. Light blue dashed line gives mean change in membrane current over time from rd10 cones in K-Asp internal (n=13).

In animals 3½ weeks old, when the eyes were large enough to make patch-clamp recording feasible, the membrane capacitance of rd10 cones was already significantly decreased. From data of all ages, rd10 cones had an average membrane capacitance of 2 ± 1 pF (n = 42) in comparison to WT cones of 6 ± 2 pF (n = 22, p<0.01), reflecting the loss of the outer segment. We could detect no change in WT membrane capacitance with animal age from linear regression (WT slope = 0.05, 95% confidence interval (CI) = [-0.06, 0.17]); however, for rd10 cones there was a small but statistically significant increase in membrane capacitance (rd10 slope = 0.14, 95% CI = [0.03, 0.26]), probably resulting from the growth of processes from the proximal end of the cell.

### Light responses of rd10 cones

We were surprised to discover that many of the cones even without outer segments were able to respond to light (Figure 1A). For those cells from which we were able to make stable recordings, light responses were observed in 4 of 4 cones at 3.5-weeks-old, 7 of 10 cones at 6-weeks-old, and 1 of 3 cones at 9-weeks-old. We may be underestimating the percentage of responding cells especially at 9 weeks, because of the increasing difficulty of achieving stable patch-clamp recordings from cones as degeneration progressed. Light responses even to our brightest stimuli were small (Figure 1A, 2 ± 1 pA, n = 10) in comparison to WT cones (Figure 1B, 27 ± 9.7 pA, n = 22). Cones were also less responsive. A precise measure of sensitivity could not be obtained, because of the difficulty of estimating the pigment concentration in cells without outer segments, and because of our inability to provide sufficiently bright light to investigate the full range of cell responsiveness with our photostimulator. We nevertheless estimate that degenerating cones were at least two orders of magnitude less responsive than WT cones [19]. We were also surprised to discover that the cones of rd10 mice maintained a normal resting membrane potential of −50 ± 9 mV (n=40), compared to −47 ± 5 mV (n=18) in WT cones (p=0.085). Additionally, linear regression showed no significant change in resting membrane potential with age for either WT or rd10 cones (WT slope = 0.51, 95% CI = [-0.46, 1.49]; rd10 slope = −0.62, 95% CI = [-2.16, 0.92]).

Since cGMP-gated channels, which are open in darkness, are largely responsible for the depolarized membrane potential of the photoreceptor, we were curious to know how abundant these channels are in degenerating cones. We therefore attempted to test more generally for the presence of cGMP-gated channels by blocking them with 500 μM L-cis diltiazem [20]. Unfortunately, we were unable to detect a consistent effect of this agent even on WT cones (not shown) [21]. As an alternative approach, we attempted to activate the channels by dialyzing the cones with 100 μM cGMP. In a WT cone, diffusion of cGMP into the outer segment produced a large inward current (Figure 1C, left) which could be suppressed by light (Figure 1C, right). As flashes were made brighter, the cGMP phosphodiesterase (PDE) was activated more strongly, and hydrolysis of cGMP became more rapid than diffusion from the whole-cell pipette into the outer segment. As a result, the membrane current returned to the level of the initial value at the beginning of the experiment, set to zero pA.

When we dialyzed rd10 cones with cGMP, we saw a similar increase in inward current. The cell in Figure 1D from a 9-week-old mouse initially displayed small responses to bright light and developed a large current as cGMP dialyzed into the cell. This current rose much faster than in a WT cone, presumably because cGMP did not have to diffuse through the cone cilium into the outer segment to reach the majority of the cGMP-gated channels. We were unable to extinguish this current even with the brightest light flash we gave (2.7×10^7^ photons μm^−2^), perhaps because the rate of entry of cGMP into the much smaller cell body of the cone greatly exceeded the ability of the cell to activate PDE. The average change in current in an rd10 cone was about half that in a WT cone (rd10, −55 ± 21 pA, n = 9; WT, −95 ± 10 pA, n = 3; p = 0.018), but the capacitance and therefore total membrane area of an rd10 cone was much smaller. The density of cGMP-gated channels may therefore be approximately the same or perhaps even greater in a degenerating cone than in a WT cone.

These observations, though indirect, suggest that the depolarized membrane potential of an rd10 cone may be produced largely by open cGMP-gated channels as in a WT cone. We were unable to substantiate the contribution of other channels with channel blocking agents: the only effect we saw was a slow depolarization of the membrane potential probably caused by gradual deterioration of the recording. For example, blocking hyperpolarization-activated, cyclic-nucleotide-gated (HCN) channels with 5 mM external Cs^+^ solution [22, 23] slowly decreased the resting potential from −52 ± 7 to −45 ± 7 (n = 5). Similar effects were observed with 25 mM external tetraetylyammonium (TEA) to block the sustained voltage-gated K^+^ channel [24] (−52 ± 19 to −40 ± 17, n = 4) and 250 μM niflumic acid to block the Ca^2+^-activated Cl^−^ channel [24] (−45 ± 6 to −39 ± 8, n = 3). None of these changes was statistically significant.

### Degenerating cones have rhodopsin and may retain organized membrane

In electron micrographs of rd10 retinas, we identified degenerating cone photoreceptors with two main types of morphology at 6 or 9 weeks of age. One type included a clearly evident cilium, with rudimentary membrane amplification. From these cells we could detect cone opsin (with a combination of blue and red/green cone opsin antibodies) in both the cilium and the inner segment (Figure. 2A and 2C). These cones were localized mostly in the periphery of the retina remote from the region where we made our recordings. The other type of cell lacked a cilium, but cone opsin was nevertheless detected throughout the inner segment (Figure. 2B and 2D). Cells with this morphology were more common in the central area of the retina. Even though cones with either morphology lacked an elaborated outer segment, they still expressed opsin, which was distributed throughout the inner segment and, when present, the cilium. With extensive searching of the periphery of the retina in the electron microscope, a very small number of cones could be observed with a stack of well-organized disk-like membranes, providing an outer segment up to half the length of a WT mouse cone outer segment. Cones of this morphology were only observed at more than 1.5 mm away from the center of the eye.

**Figure 2.**
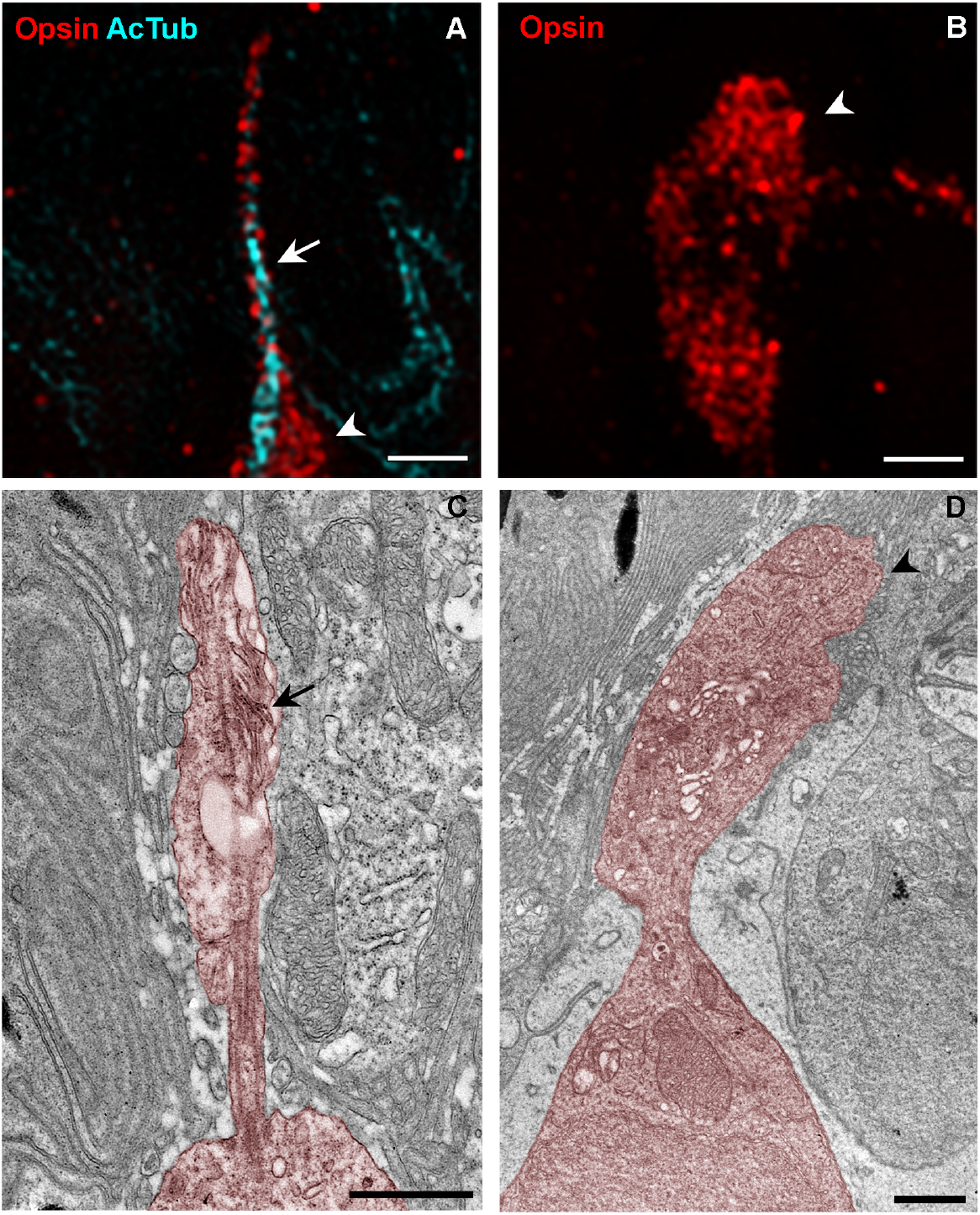
Morphology of cone photoreceptors in 6-9 week old rd10 mice from super-resolution SIM imaging of immunofluorescence and from electron microscopy. (A and B) SIM images. (A) Ciliated cone photoreceptor cell. The cilium (arrow) is immunolabeled with acetylated tubulin (cyan). Inner segment (arrowhead) as well as the cilium is labeled with cone opsin antibodies (red). (B) Cone photoreceptor cell lacking a cilium. Inner segment only (arrowhead) is labeled by cone opsin antibodies (red). (C and D) Electron micrographs, a single cone photoreceptor has been colorized. (C) Ciliated cone photoreceptor with residual membrane amplifications (arrow). (D) Cone photoreceptor with inner segment (arrowhead) but lacking cilium. Scale bars (A) – (D), 1 μm.

### Ca^2+^ channels and other voltage-dependent currents

The greatest contributor to turnover of ATP in cones is the energy required to pump out Ca^2+^ entering through voltage-gated Ca^2+^ channels at the synaptic terminal [12]. Degenerating cones lack synaptic pedicles, and we had expected that they would also have much smaller, if any voltage-gated Ca^2+^ currents. We were therefore surprised to discover that cones in rd10 retinas had a robust Ca^2+^ current with an average peak value of −29 ± 10 pA (n = 19), not significantly different from WT cones (−23 ± 9 pA, n = 9, p = 0.13). Additionally, there was no significant difference in the voltage sensitivity of the channel, with peak current in rd10 cones seen at an average membrane voltage of −40 ± 5 mV (n = 19) comparted to WT cones at −36 ± 10 mV (n = 9, p = 0.28). When the peak current of rd10 cones was plotted as a function of age, linear regression showed no significant change with time (slope = −1.15, 95% CI [-3.51, 1.21]). We were also unable to detect differences in the amplitude or voltage dependence of the HCN current between WT and rd10 cones (Figure 3B). At a voltage of −105 mV, for example, mean values for WT were −56 ± 20 pA (n = 8), and for rd10 − 74 ± 34 pA (n = 23, p = 0.12). Linear regression showed no significant change in the HCN current with age for either WT or rd10 cones (WT slope = −2.17, 95% CI = [-6.00,1.66]; rd10 slope = −2.12, 95% CI = [-10.51, 6.11]).

**Figure 3.**
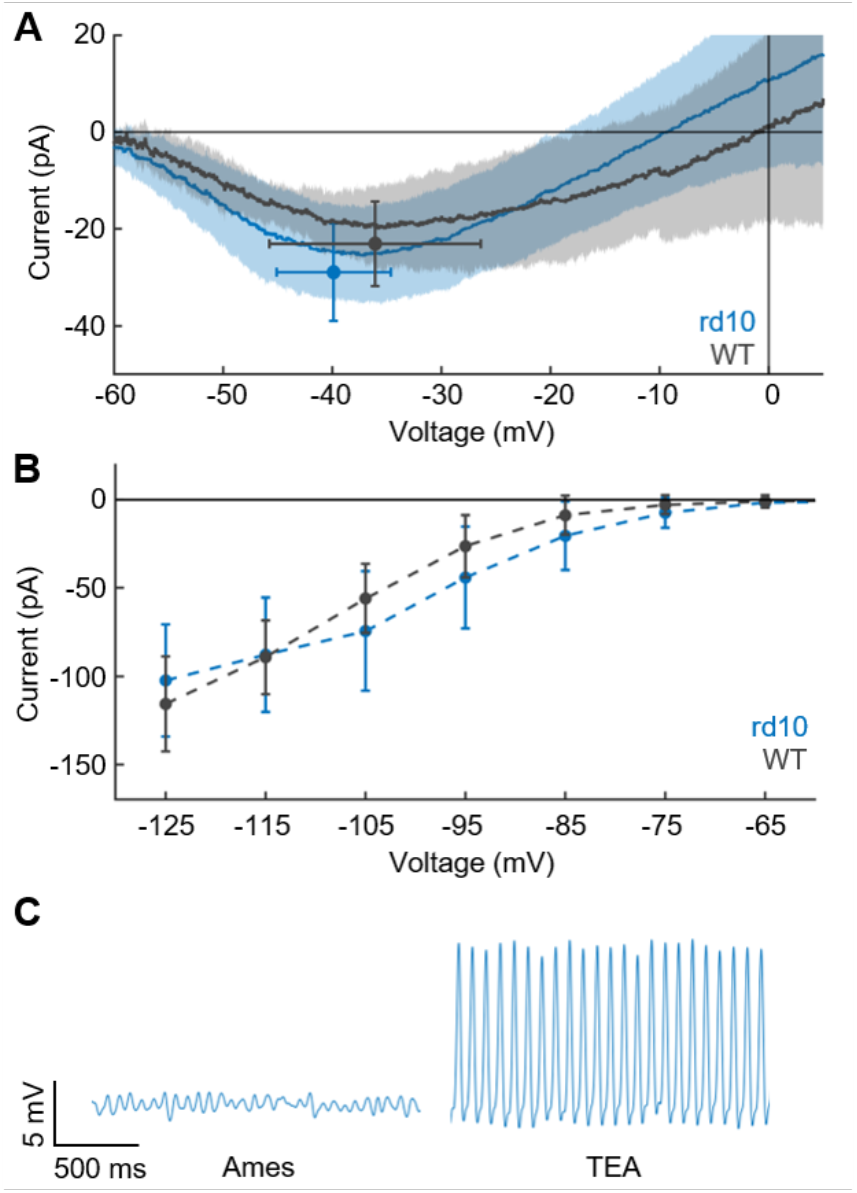
Membrane currents in degenerating cones. (A) Voltage-gated Ca^+2^ current recorded with ramp protocol and Cs-TEA internal solution to block voltage-gated K^+^ channels (see Methods). Average current is plotted as a function of membrane voltage for WT cones (gray, n = 9) and rd10 cones (blue, n = 19). Shaded area is SD. Points are average peak current with SD. (B) HCN-channel current (i_h_) recorded from holding potential of −50mV with 300-ms steps [24] to voltages of −55 to −125mV in increments of 10mV; responses are mean currents during the last 50 ms of the voltage pulse. Steady-state current is plotted versus membrane potential for WT (gray, n = 8) and rd10 (blue, n = 23) cones; bars are SD. (C) Current-clamp recordings from 5-week-old rd10 animal. Membrane voltage showed small oscillations in Ames’ media (left); washing on 25mM external TEA triggered large spikes (right) probably produced by voltage-gated Ca^2+^ conductance, indicating presence of sustained, TEA-sensitive voltage-gated K^+^ channels [24].

In many of the rd10 cones (but none of the WT cones), we observed spontaneous oscillations of membrane potential or even action potentials. An example is given in Figure 3C (left). When for 4 cells showing small oscillations we perfused the retinal slice with 25 mM TEA, they all began to show larger action potentials (Figure 3C, right), as previously observed for amphibian rods in TEA [25, 26]. Although the ion dependence of these action potentials was not investigated, it is likely that they are produced by voltage-gated Ca^2+^ channels. The effect of TEA indicates that rd10 cones have a sustained voltage-gated K^+^ conductance like that in WT cones which stabilizes the membrane potential in darkness [24], though we made no attempt to measure its amplitude.

### Light responses of second-order cells

We wondered whether the small light responses in cones were able to drive changes in membrane potential or current in second-order horizontal and bipolar cells. We focused on retinas of 8 – 9 week-old mice, whose outer nuclear layer in most of the retina had almost entirely degenerated and whose cones—when present—lacked outer segments and synaptic pedicles but often had small processes extending from the proximal end of the photoreceptor. The recordings in Figure 4 are representative of 10 ON bipolar cells from four mice (A), 5 OFF bipolar cells from 2 mice (B), and 3 horizontal cells from 3 mice (C), with mean maximum response amplitudes from all of the cells of −22 pA (A), 85 pA (B), and 97 pA (C). Responses were quite variable, with ranges of −3 to −52 pA for ON bipolar cells, 12 to 280 pA for OFF bipolar cells, and 34 to 240 pA for horizontal cells. Responses were usually smaller in amplitude than for WT cones but not always (Figure 4B). Sensitivity was again difficult to estimate, but half-maximal responses required light intensities between 10 and 100 times greater than for normal cones [19].

**Figure 4.**
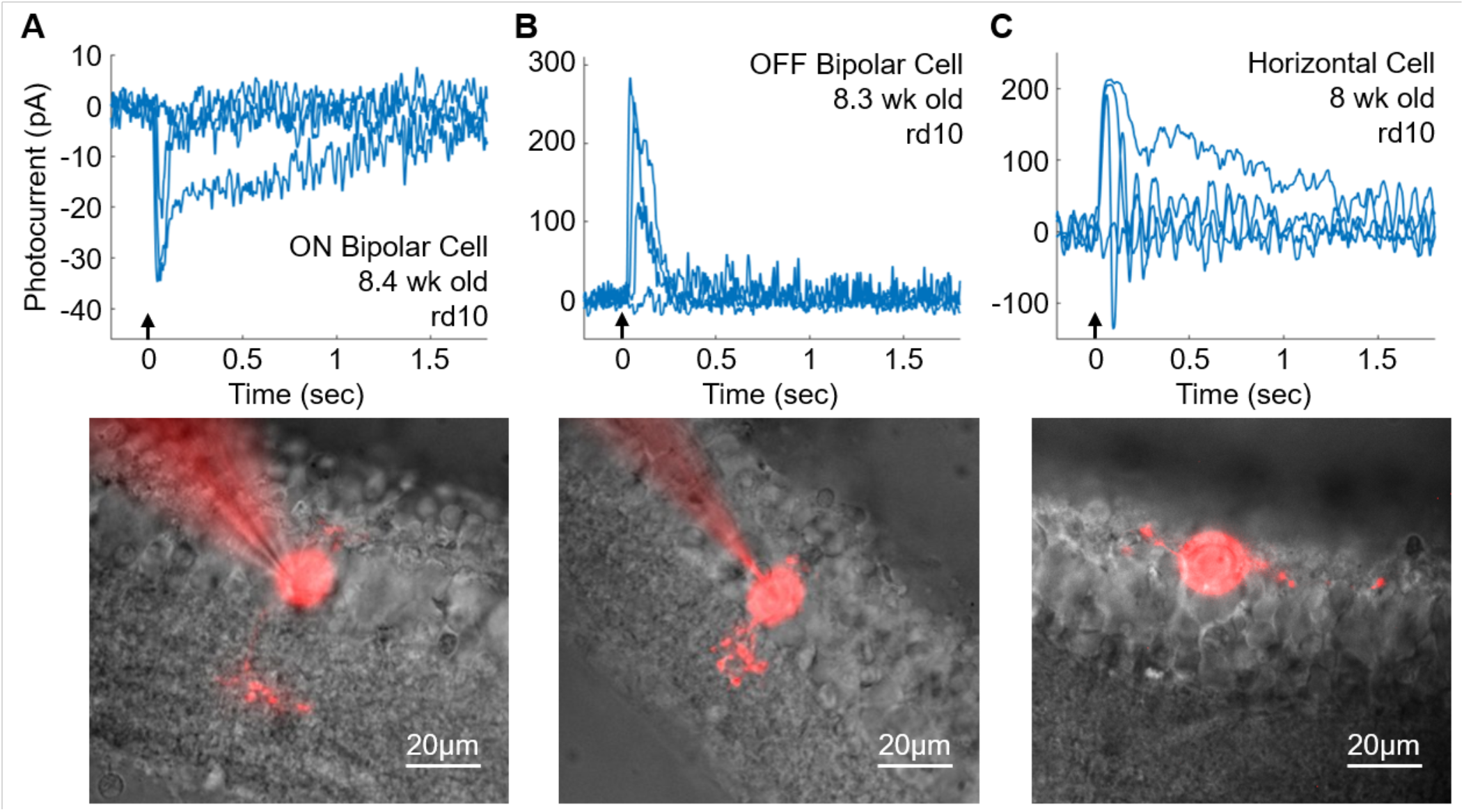
Light-evoked responses in second-order retinal neurons from 8-9 week-old rd10 retinas. (A), ON Bipolar Cell; (B), OFF bipolar cell; (C), horizonal cell. Stimuli were 10ms flashes at 405nm (arrow); flash intensities were 6.1×10^1^, 2.4×10^3^, 2.6×10^4^, and 1.5×10^7^ photons μm^−2^ in A; 2.4×10^3^, 2.6×10^4^, 2.6×10^5^ and 5.4×10^7^ photons μm^−2^ in B; and 2.6×10^5^, 8.7×10^5^, 9.1×10^6^, and 3.0×10^7^ photons μm^−2^ in C. Below: Cells of A-C filled with Alexa-750; note that outer nuclear layer has almost entirely degenerated in all three images. Scale bars, 20μm.

The morphologies of the recorded cells filled with Alexa-750 are shown below the recordings. The bipolar cells extended dendritic processes up towards the degenerated outer nuclear layer. Their axonal processes could be seen reaching down into the inner plexiform layer. The horizontal cell in Figure 4C was sitting on the outer edge of the degenerated retina, with long dendritic arbors extending laterally.

### Retinal output is robust despite deteriorating cone morphology

Finally, we examined how the degeneration o f photoreceptors of eight-week-old rd10 mice impacts the signals sent from the retina to the rest of the brain. We began by measuring the fidelity of visual signaling by recording retinal ganglion cell (RGC) responses with a large-scale multi-electrode array [27] while presenting a natural movie. Specifically, we measured the mutual information between the stimulus and the responses [28]. Under scotopic (∼1 Rh*/rod/s) and mesopic (∼100 Rh*/rod/s) conditions, the RGC spike trains conveyed substantially less information about the natural movie (Figure 5A-D), presumably because of the massive decrease in the number of rods in 8-week-old animals. Surprisingly, under photopic (∼10,000 Rh*/rod/s) conditions, despite extensive changes in cone morphology and in the amplitude of their light responses, information rates among RGCs were remarkably similar to those in healthy retinas (Figure 5E-F).

**Figure 5.**
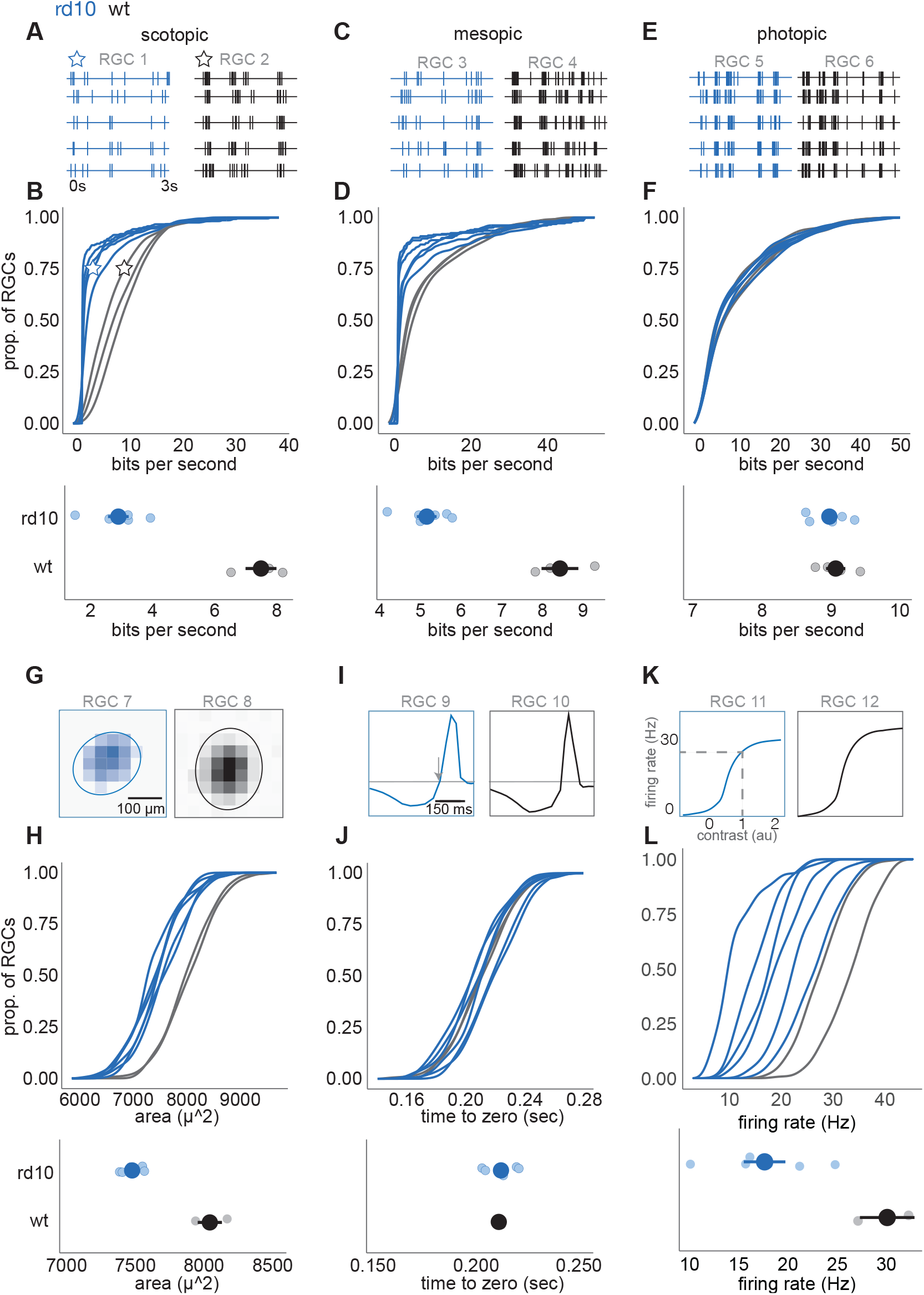
Retinal ganglion cell responses at photopic light level are similar to wild-type in 2 month rd10 retinas. (A) Spike times from two example RGCs to 5 trials of a natural movie at a scotopic light level (1 R*/rod/s) RGC 1 (blue) is rd10, RGC 2 (black) is WT. (B) Top: cumulative distributions of information rates across RGCs, each line is a different retina. Blue is rd10, black is WT. Stars indicate the location of example RGCs 1 and 2 in A. Example RGCs are near the 75^th^ percentile information rate. Bottom: mean ± 2 S.E. information rates between rd10 and WT retinas at the scotopic light level (large points are means across under photopic conditions between WT and eight-week-old rd10 mice was a 39% decrease in response retinas, small points are individual retinas). (C and E) Same as A, but measured at mesopic (100 Rh*/rod/s) and photopic (10,000 Rh*/rod/s) light levels, respectively. (D and F). Same as B, but measured at mesopic and photopic light levels. (G) Example spatial receptive fields from two RGCs at the photopic light level. RGC 7 (blue) is rd10, RGC 8 (black) is WT. Example cells lie near the 75^th^ percentile of receptive field area. (H) Top: cumulative distribution of receptive field areas over RGCs for each retina. Blue is rd10, black is WT. Bottom: mean ± 2 SE of receptive field areas (large points are means across retinas, small points are individual retinas). (I) Example temporal receptive fields from two RGCs near the 75^th^ percentile for the time-to-zero crossing. Arrow indicates time-to-zero crossing. RGC 9 is rd10, RGC 10 is WT. (J) Same as H, but for the time-to-zero crossing of the temporal receptive fields. (K) Example contrast response functions from two RGCs near the 75^th^ percentile for gain. Dashed lines indicate spike rate at a generator signal equal to one, which was used to estimate response gain. (L) same as H, but for gain of the contrast response functions and gain estimates.

These measurements capture the fidelity of visual signaling (the extent of reproducibility from one trial to the next) but do not indicate what is being signaled. Though information rates remain high under photopic conditions, the RGCs could conceivably exhibit substantial changes in receptive field structure, indicating a change in their feature selectivity. To test for this possibility, we focused on the photopic condition and used checkerboard noise to estimate the spatial and temporal receptive fields of RGCs [27, 29]. Spatial receptive fields were systematically smaller in rd10 mice eight weeks of age (Figure 5G-H) but only by about 7% (p = 0.02). The mean duration of temporal integration was not significantly different between WT and eight-week-old rd10 retinas (Figure 5I-J). Finally, we analyzed contrast-response functions (see Methods) of WT and rd10 RGCs (Figure 5K-L). The contrast-response function identifies the mean spike rate produced for a given similarity between a stimulus and the spatiotemporal receptive field. This similarity is frequently referred to as the generator signal, which is computed as the convolution between the stimulus and the spatiotemporal receptive field. RGCs with higher gain will produce more spikes in response to stimuli with a given similarity between the receptive field and the stimulus. WT RGCs as a population exhibited higher gain than rd10 RGCs (Figure 5K-L). For example, for WT RGCs the output produced by a generator signal of one was 30.5 ± 4.2 spike/s, while for rd10 RGCs the output was 18.6 ± 1.4 spikes/s (mean ± S.E., p = 0.0001). Thus, the most significant change in RGC signaling gain accompanied by a 7% decrease in RF size, while temporal integration and information rates were indistinguishable.

## DISCUSSION

We have made the discovery that degenerating cones, even lacking outer segments and synaptic pedicles, still respond to light and mediate surprisingly robust processing of visual signals. Although most cones will eventually die [15-17] and the animals become blind, our results indicate that for a period of many weeks in the mouse retina while degeneration is occurring, cones and the retina can still respond to light [14]. These results indicate that efforts to revive degenerating cones—provided they are undertaken sufficiently early—have the potential to restore substantial visual function.

Our discovery that cones maintain a depolarized resting membrane potential confirms previous observations [4] and is consistent with the continued presence of cyclic-nucleotide-gated channels in the degenerating cells (Figure 1D). We cannot be sure that all of these channels are functional, since cone inner segments are known to have cGMP-gated channels [30] not all of which participate in phototransduction [31]. We have no information about the resting concentration of cGMP, and inner-segment channels may be positioned too far away from the transducing membrane of the inner segment to produce a light response (see Figure 2). We have nevertheless shown that at 8 – 9 weeks of age, light responses can be recorded from second-order bipolar and horizontal cells in retinal slices and from ganglion cells in a retinal whole mount with a multi-electrode array. Light responses in retinal neurons could only be generated if cones were sufficiently depolarized to have open voltage-gated Ca^2+^ channels in darkness. This depolarization could conceivably be produced by leak across the cone membrane due to injury during degeneration or recording, but the global mean value of the input resistance we measured from rd10 cones during our experiments (1800 ± 320 MΩ, n = 42) is about a factor of 2 greater than that of normal cones (890 ± 190 MΩ, n = 22). This difference probably reflects the decrease in membrane area and resting conductance resulting from loss of the outer segment. We have no indication that the degenerating cones in our recordings were leaky or otherwise impaired. The simplest explanation of our results is that degenerating cones are behaving similarly to normal cones with open cyclic-nucleotide-gated channels in darkness, despite the marked change in their morphology and reduced light sensitivity.

Perhaps the most unexpected result from this study comes from the RGC measurements in two-month-old rd10 mice, when nearly all of the rods are gone, and the outer nuclear layer consists in most of the retina of only a single layer of cells nearly all of which are cones [15]. Even though cones have undergone extensive changes in morphology and no longer have a recognizable outer segment (Figure 2), their signals still propagate to second-order cells (Figure 4) as well to ganglion cells (Figure 5). The visual signals of RGC are surprisingly robust, exhibiting small decreases in receptive-field size (Figure 5G-H) and sensitivity (Figure 5K-L) but relatively normal temporal integration (Figure 5I-J) and information rates (Figure 5E-F). These results indicate that most of the mechanisms of visual integration in the outer and inner plexiform layers are intact and continue to function, providing the mouse—at least under photopic conditions—with visual function as nearly normal as possible. Though ganglion-cell responses in rd10 mice eventually degrade and disappear [32], the animals seem nevertheless to preserve substantial visual function for a prolonged period.

The RGC results in rd10 mice are similar to recent measurements in a different model of RP, the *Cngb1*^*neo/neo*^ mouse [33]. In this channelopathy, rods degenerate but much more slowly and are not lost until about 6 months of age. RGCs in *Cngb1*^*neo/neo*^ also retain surprisingly normal receptive field structure and information rates at cone-mediated light levels despite extensive changes in cone morphology and the loss of outer segments [14]. Thus, the preservation of cone-mediated signaling pathways may be a general feature in RP, which spans different underlying causes of rod death and different rates of degeneration.

Many approaches are presently being tested that use gene therapy to ameliorate genetically inherited eye disease [see for example 34]. The efficacy of these approaches has been questioned because synaptic connections of photoreceptors and other retinal neurons have been shown to undergo considerable remodeling during degeneration [see for example 35]. This remodeling could conceivably prevent recovery of normal vision even after restoration of photoreceptor function. Our results show, in contrast, that the retina has a surprising ability to continue to detect and process the visual scene even as degeneration is occurring, in spite of (or perhaps because of) synapse remodeling. Our observations give new hope that strategies to recover cone photoreceptor function may be able to provide substantial improvement in visual function, as long as some photoreceptors are still alive and can be rescued.

## ACKNOWLEDGEMENTS

We thank Rikard Frederiksen for helpful discussion and Ekaterina Bikovtseva for technical assistance. This work was supported by NEI R01 EY033035 and NEI R01 EY027442 to D.S.W.; NEI R01 EY27193 to G.D.F.; NEI R01 EY001844 to G.L.F.; NEI R01 EY27193 and EY29817 to A.P.S.; a fellowship of the UCLA EyeSTAR program of the UCLA Department of Ophthalmology to E.M.E.; a BrightFocus Foundation Postdoctoral Fellowship to A.E.P.; an unrestricted grant from Research to Prevent Blindness USA to the UCLA Department of Ophthalmology; and NEI Core Grant (P30) EY00311 to the Jules Stein Eye Institute.

## AUTHOR CONTRIBUTIONS

Conceptualization: G.L.F., A.P.S., E.M.E, M.S., G.D.F.

Data Collection: E.M.E., A.E.P., J.R., Y.J.

Formal Analysis: E.M.E., A.E.P., M.T.

Writing Original Draft: E.M.E., A.E.P., G.D.F., G.L.F.

Writing Reviewing and Editing: G.L.F., D.S.W., G.D.F., A.P.S.

Project Administration: M.S., D.S.W., G.D.F., G.L.F., A.P.S.

## DECLARATION OF INTERESTS

The authors declare no competing interests.

## CONTACT FOR REAGENT AND RESOURCE SHARING

Further information and requests for resources and reagents should be directed to and will be fulfilled by the lead contact, Alapakkam P. Sampath.

## METHODS

### Animal use and care statement

This study was carried out in accordance with the recommendations in the Guide for the Care and Use of Laboratory Animals of the National Institutes of Health, and the Association for Research in Vision and Ophthalmology Statement for the Use of Animals in Ophthalmic and Vision Research. The animal use protocol for cones and retinal cells was approved by the University of California, Los Angeles Animal Research Committee. All procedures for ganglion-cell recordings were approved by the Duke University Institutional Animal Care and Use Committee.

### Solutions

After retinal dissection, retinal tissue was kept alive in a specially formulated Ames’ medium (pH 7.4, osmolarity 284 mOsm), either buffered with bicarbonate (Ames’-Bicarb) and bubbled with carbogen gas (95% O_2_, 5% CO_2_); or, for embedding and slicing, buffered with HEPES (Ames’-HEPES) and bubbled with 100% oxygen. For certain experiments, channel-blocking agents were added to the Ames’-bicarb bath solution as follows: isradipine (ISR, 10 μM), to block L-type voltage-gated Ca^2+^ channels; niflumic acid (NFA, 250 μM), to block Ca^2+^-activated Cl^−^ channels; tetraethylammonium (TEA, 25 mM), to block sustained voltage-gated K^+^ channels; cesium (Cs^+^, 5mM), to block HCN channels; and L-cis-diltiazem (DILT, 500 μM), to block CNG channels. The standard internal solution for whole-cell patch clamp recordings was a potassium aspartate solution (K-Asp) containing (in mM): 125 potassium aspartate, 10 KCl, 10 HEPES, 5 *N*-methyl-D-glucamine-HEDTA, 0.5 CaCl_2_, 0.5 MgCl_2_, 0.1 ATP-Mg, 0.5 GTP-Tris, 2.5 NADPH (pH 7.3, 280 mOsm). For experiments investigating the presence of CNG channels, cGMP (50-100 μM) was added to the standard internal solution. In other experiments, a cesium-TEA (Cs-TEA) based internal solution was used to block K^+^ conductances. The Cs-TEA internal solution contained (in mM) 110 CsCH_3_O_3_S, 12 TEA-Cl, 10 HEPES, 10 EGTA, 2 QX-314-Br, 11 ATP-Mg, 0.5 GTP-Tris, 0.5 MgCl_2_, 1 NAD^+^ (pH 7.3, 280 mOsm). QX-314 is 2-[(2,6-dimethylphenyl) amino]-N,N,N-triethyl-2-oxoethanaminium; it is a derivative of the anesthetic lidocaine and is a blocker of voltage-gated Na^+^ channels.

### Preparation of retinal slices

Prior to each experiment, animals were dark adapted overnight. The retinal dissection and preparation of retinal slices were performed under infrared illumination. Animals were euthanized by cervical dislocation. The dorsal aspect of each eye was marked prior to enucleation. After enucleation, the globes were moved to an Ames’-bicarb bath. The anterior segment was carefully removed, including cornea and lens. The posterior eye cup was bisected into dorsal and ventral halves. From one half, a rectangular section of retina was cut from the central region of the retina and then carefully separated from the retinal pigment epithelium. The isolated section of retina was embedded in 3% low-temperature-gelling agar dissolved in Ames’-HEPES solution. The retina was sliced in cold Ames’-HEPES with a vibratome to obtain 200 μm-thick slices of retina. Slices were made perpendicular to the retina to obtain a cross-section of retinal tissue with intact retinal circuitry. The retinal slice selected for recording was mounted on a recording dish; the slice was held in place by a custom-made anchor and moved to the microscope. The remaining retinal tissue and retinal slices were stored in a light-tight container with Ames’-bicarb solution and kept at 32ºC. The tissue under the microscope was perfused with Ames’-bicarb solution and kept at 35ºC.

### Identifying cones

In WT tissue, cones in slices can be distinguished from rods by their characteristic morphological and somatic features [19]. Identifying cones in the degenerated tissue was more challenging since many of the rod somata are swollen and have disrupted nuclear architecture. Additionally, it was difficult to remove cell layers from the ONL of the degenerating tissue with a vacuum pipette, as is normally done in WT tissue to expose healthy cell bodies deeper in the retinal slice. Cones could however be found in the degenerating tissue along the top edge of the outer nuclear layer and generally had a small though easily recognized inner segment. Targeting the inner segment was the most reliable way to make patch-clamp recordings from these cells.

### Whole-cell patch-clamp recordings

Whole-cell patch-clamp recordings were performed with borosilicate glass micropipettes (15-19 MOhm) filled with a K-Asp internal solution. Different internal solutions were used for specific experiments, as noted below. Cells were recorded in the voltage-clamp mode and initially held at −50mV. After breaking into the cell, recordings could be made in voltage-clamp or switched to current-clamp mode. Series resistance was compensated at 75-80%. Light stimuli (10-20ms) were delivered from a monochromatic 405nm LED, at a wavelength that is approximately the isosbestic point of the S and M mouse cone opsins and stimulates both opsins with nearly equal efficiency. Light intensities are given in photons μm^−2^ rather than in light-activated cone visual pigment molecules (P*). Conversion to P* requires knowledge of the cone outer-segment collecting area, and degenerating cones lack an outer segment. For most experiments, recording pipettes included a fluorescent dye, either Alexa-750 or Alexa-647, so that cellular morphology could be imaged after electrophysiological recordings were completed. Electrophysiology data were filtered at 500 Hz, sampled at 10kHz, and acquired with Symphony (https://open-ephys.org/symphony/), an open-source MATLAB-based data-acquisition system.

To record the voltage-gated Ca^2+^ current, we used a Cs-TEA internal solution to block K^+^ currents including i_h_ and one of two ramp protocols. In both protocols, cells were initially held at −50mV and were then either stepped to −70mV and held there for 200 - 500 ms, to allow all the Ca^2+^ channels to close before ramping the voltage to +20 mV; or were stepped to −100mV and immediately ramp to +30 mV. Ramp rate was 80 mV/sec. Leak was subtracted from a section of the trace prior to the opening of the Ca^2+^ channels, where the change in current was linear and proportional to the leak voltage. Data from the two protocols were similar and were combined for population analysis.To characterize the HCN channel conductance, recordings were made with the K-Asp pipette solution and in Ames’ medium, without the addition of any blockers. Currents were evoked with 300 ms hyperpolarizing voltage steps from a holding potential of − 50 mV, and the value of the current response was estimated from the mean during the last 50 ms of the voltage pulse.

### Immunostaining and structured-illumination microscopy imaging

After euthanasia, retinas were removed and fixed with 4% formaldehyde in PBS (pH 7.4) for 30 minutes at 4 °C. They were then washed with PBS, blocked with 5% donkey serum in PBS-0.2%TritonX100 (v/v) for 1h at room temperature (RT), and incubated with a mixture of 1:1000 rabbit anti-blue opsin antibody (AB5407, Millipore Sigma), 1:1000 rabbit anti-red/green opsin antibody (AB5405, Millipore Sigma), YF594-conjugated acetylated tubulin antibody (YF594-66200, Proteintech), and 2% donkey serum in PBS-0.2%TritonX100 (v/v) overnight at 4 °C. After three washes of 10 minutes each at room temperature with PBS-0.2%TritonX100 (v/v), retinas were incubated with 1:750 anti-rabbit Alexa Fluor 488 antibody (ThermoFisher Scientific) and 2% donkey serum in PBS-0.2%TritonX100 (v/v) for 1h at room temperature, and washed again in PBS-0.2%TritonX100 (v/v). Whole retinas were mounted on a slide with Fluoro-Gel (17985-10, EMS), and we imaged cone photoreceptors with three-dimensional structured illumination microscopy (SIM), using a GE OMX SR microscope with a Blaze SIM module, 60× 1.42 NA objective lens and pco.edge sCMOS cameras. Z-planes were obtained 125 nm apart; we collected 15 raw images per plane (three angles and five phases). Reconstructions were performed with a Wiener filter setting of 0.001, along with channel-specific optical transfer functions in the softWoRx software package (Applied Precision) and the maximum projection of several Z-planes obtained with ImageJ software. Minor contrast and brightness adjustments were applied evenly to entire images with Adobe Photoshop software.

### Electron microscopy

After euthanasia, enucleated eyes were fixed with 2% formaldehyde and 2% glutaraldehyde in 0.1 M cacodylate buffer (pH 7.4) for 24h at 4 °C. They were then postfixed with 1% OsO_4_ (w/v) and 1 % potassium ferricyanide (w/v) for 1h. After a wash with distilled water, the samples were exposed to a graded ethanol series for dehydration, then propylene oxide, followed by resin infiltration. Samples were embedded in Araldite 502 resin (Electron Microscopy Sciences), and ultrathin sections (70 nm) were stained with uranyl acetate and lead citrate. Micrographs were acquired with a JEM-1400 Plus (JEOL) electron microscope, equipped with an Orius SC1000A (Gatan) camera and Gatan Microscopy Suite Software. Minor contrast and brightness adjustments were applied evenly to entire images with Adobe Photoshop software.

### Multi-electrode-Array Ganglion-cell recordings

Mice in their home cages were placed in a light shielded container fitted with an air pump to dark adapt overnight. Dissections took place under infrared illumination with infrared converters. Mice were decapitated, and enucleated eyes were placed in oxygenated room-temperature Ames’ solution. Eyes were hemisected and the vitreous was removed as described previously [36]. Following the vitrectomy, the retina was isolated from the retinal pigment epithelium, and a dorsal sample of retina (∼1.25mm x 1.25mm in area and centered ∼1.5 mm from the optic nerve) was placed RGC side down on an electrode array consisting of 519 electrodes with a 30 µm pitch [37, 38]. Bicarbonate-buffered Ames’ media was heated to 32ºC, and the retina was perfused at a rate of 6-8 mL/min.

### Spike Sorting

Spike sorting was performed on the raw voltage traces collected from the multi-electrode array with custom software, then manual curation [39, 40]. Events that crossed a threshold of 4-standard deviations from the mean voltage were identified as spike events. These events were accumulated by taking the electrical signal 0.5 ms preceding and 1.5 ms following the time of threshold crossing. Principal component analysis was used to reduce the dimensionality of the spike waveforms from 40 samples to five. This procedure was followed by a water-filling algorithm and expectation maximization to initialize and fit a mixture of Gaussian model for clustering spikes by their shape [38]. We retained potential cells that had a spike rate of >0.1Hz and <10% contamination, as estimated from refractory period violations.

### Visual Stimuli

Visual stimuli were presented as previously described [14]. Briefly, A gamma-calibrated OLED (Emagin, SVGA + XL Rev3) was used to focus an image onto photoreceptors with a 4x objective (Nikon, CFI Super Fluor) attached to an inverted microscope (Nikon, Ti-E). Visual stimuli were presented at three light levels, with a 5-min adaptation period between light levels of ∼1 Rh*/rod/s (scotopic), ∼100 Rh*/rod/s (mesopic), and ∼10,000 Rh*/rod/s (photopic). At each light level, checkerboard stimuli were presented for 30 minutes to calculate spatiotemporal receptive fields and contrast response functions. For scotopic and mesopic stimuli, each square of the checkerboard stimulus was 150×150 μm and refreshed at 15 Hz. For photopic stimuli, each square was 75×75 μm and refreshed at 60 Hz. Additionally, ten-second clips of natural movies [adapted from 41] were repeated one hundred times each at all light levels to calculate mutual information.

### Receptive Field Measurements

The spike-triggered average was used to estimate the linear receptive fields for RGCs in response to checkerboard noise [29]. As described previously [14], RGCs with space-time separable receptive fields were selected for this analysis, because a unique spatial and temporal filter could be identified for each cell. Space-time separable receptive fields were defined as those in which >60% of the variance in the spike-triggered average was explained by a rank-one factorization with singular value decomposition. Across three control retinas, 75% percent of RGCs met this criterion; across six 2-month-old rd10 retinas, 55% of RGCs met this criterion. To estimate the time-to-zero crossing in the temporal receptive fields, cells with biphasic filters were used, because they exhibited a well-defined zero crossing time. 70% of space-time separable spike-triggered averages met this criterion. The contrast response function for each RGC was given by the static nonlinearity from a linear-nonlinear model fit to each RGC response [29]. Response gain was estimated as the spike rate produced for a stimulus that produced a ‘generator signal’ equal to one. The generator signal was calculated as the convolution between the spatiotemporal receptive field and the checkerboard noise across all frames of the stimulus. To equate receptive fields and the effectiveness of the stimulus across cells, the spatiotemporal temporal RF estimated from the spike-triggered average was normalized to have a vector length equal to one.

### Quantification and statistical analyses

#### Cones

Cone data were analyzed with custom scripts written in MATLAB. All averages are reported as mean ± standard deviation. Comparisons between means of WT control data and rd10 data were performed with the Wilcoxon Rank Sum test, a non-parametric comparison that does not assume equal variance. A p-value of less than 0.05 was considered significant. To analyze changes in parameters over time, data were fit with linear regression without fixing the y-intercept, with 95% confidence intervals (CI) reported for the regression coefficients. Fits in which the 95% CI for the slope included zero were considered to indicate that there was no significant change of the parameter over time.

#### Information Analysis for Ganglion Cells

To measure the reliability with which RGCs responded to natural movies, we estimated the mutual information between the spiking responses and the stimulus with the ‘direct method’ described previously [14, 42]. Spike trains were binned according to entropy estimates that achieved the Ma Upper Bound [43] and ranged from bins of 4-6 milliseconds and formed patterns of 3-6 bins. Mutual information was thus computed as:

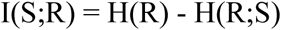

where H(R) is the entropy in the response, and H(R;S) is the entropy of the response conditioned on the stimulus.

## Notes

### Competing Interest Statement

The authors have declared no competing interest.

## REFERENCES

1. Daiger, S.P., Bowne, S.J., and Sullivan, L.S. (2015). Genes and mutations causing autosomal dominant retinitis pigmentosa. Cold Spring Harbor Perspectives in Medicine 5, a017129.

2. Punzo, C., Kornacker, K., and Cepko, C.L. (2009). Stimulation of the insulin/mTOR pathway delays cone death in a mouse model of retinitis pigmentosa. Nat Neurosci 12, 44–52.

3. Barone, I., Novelli, E., and Strettoi, E. (2014). Long-term preservation of cone photoreceptors and visual acuity in rd10 mutant mice exposed to continuous environmental enrichment. Mol Vis 20, 1545–1556.

4. Busskamp, V., Duebel, J., Balya, D., Fradot, M., Viney, T.J., Siegert, S., Groner, A.C., Cabuy, E., Forster, V., Seeliger, M., et al. (2010). Genetic reactivation of cone photoreceptors restores visual responses in retinitis pigmentosa. Science 329, 413–417.

5. Lin, B., Masland, R.H., and Strettoi, E. (2009). Remodeling of cone photoreceptor cells after rod degeneration in rd mice. Exp Eye Res 88, 589–599.

6. Amamoto, R., Wallick, G.K., and Cepko, C.L. (2022). Retinoic acid signaling mediates peripheral cone photoreceptor survival in a mouse model of retina degeneration. eLife 11.

7. Wang, W., Lee, S.J., Scott, P.A., Lu, X., Emery, D., Liu, Y., Ezashi, T., Roberts, M.R., Ross, J.W., Kaplan, H.J., et al. (2016). Two-Step Reactivation of Dormant Cones in Retinitis Pigmentosa. Cell Rep 15, 372–385.

8. Wong, F., and Kwok, S.Y. (2016). The Survival of Cone Photoreceptors in Retinitis Pigmentosa. JAMA Ophthalmol 134, 249–250.

9. Venkatesh, A., Ma, S., Le, Y.Z., Hall, M.N., Ruegg, M.A., and Punzo, C. (2015). Activated mTORC1 promotes long-term cone survival in retinitis pigmentosa mice. J Clin Invest 125, 1446–1458.

10. Léveillard, T., Mohand-Said, S., Lorentz, O., Hicks, D., Fintz, A.C., Clerin, E., Simonutti, M., Forster, V., Cavusoglu, N., Chalmel, F., et al. (2004). Identification and characterization of rod-derived cone viability factor. Nat Genet 36, 755–759.

11. Ait-Ali, N., Fridlich, R., Millet-Puel, G., Clerin, E., Delalande, F., Jaillard, C., Blond, F., Perrocheau, L., Reichman, S., Byrne, L.C., et al. (2015). Rod-derived cone viability factor promotes cone survival by stimulating aerobic glycolysis. Cell 161, 817–832.

12. Ingram, N.T., Fain, G.L., and Sampath, A.P. (2020). Elevated Energy Requirement of Cone Photoreceptors. Proceedings National Academy of Science 117, 19599–19603.

13. Okada, T., Ernst, O.P., Palczewski, K., and Hofmann, K.P. (2001). Activation of rhodopsin: new insights from structural and biochemical studies. Trends Biochem Sci 26, 318–324.

14. Scalabrino, M.L., Thapa, M., Chew, L.A., Zhang, E., Xu, J., Sampath, A.P., Chen, J., and Field, G.D. (2022). Robust cone-mediated signaling persists late into rod photoreceptor degeneration. eLife 11.

15. Barhoum, R., Martinez-Navarrete, G., Corrochano, S., Germain, F., Fernandez-Sanchez, L., de la Rosa, E.J., de la Villa, P., and Cuenca, N. (2008). Functional and structural modifications during retinal degeneration in the rd10 mouse. Neuroscience 155, 698–713.

16. Chang, B., Hawes, N.L., Pardue, M.T., German, A.M., Hurd, R.E., Davisson, M.T., Nusinowitz, S., Rengarajan, K., Boyd, A.P., Sidney, S.S., et al. (2007). Two mouse retinal degenerations caused by missense mutations in the beta-subunit of rod cGMP phosphodiesterase gene. Vision Res 47, 624–633.

17. Gargini, C., Terzibasi, E., Mazzoni, F., and Strettoi, E. (2007). Retinal organization in the retinal degeneration 10 (rd10) mutant mouse: a morphological and ERG study. J Comp Neurol 500, 222–238.

18. Wang, T., Reingruber, J., Woodruff, M.L., Majumder, A., Camarena, A., Artemyev, N.O., Fain, G.L., and Chen, J. (2018). The PDE6 mutation in the rd10 retinal degeneration mouse model causes protein mislocalization and instability and promotes cell death through increased ion influx. J Biol Chem 293, 15332–15346.

19. Ingram, N.T., Sampath, A.P., and Fain, G.L. (2019). Voltage-clamp recordings of light responses from wild-type and mutant mouse cone photoreceptors. Journal of General Physiology 151, 1287–1299.

20. Haynes, L.W. (1992). Block of the cyclic GMP-gated channel of vertebrate rod and cone photoreceptors by l-cis-diltiazem. J Gen Physiol 100, 783–801.

21. Xu, H., Jin, N., Chuang, J.Z., Zhang, Z., Zhong, X., Zhang, Z., Sung, C.H., Ribelayga, C.P., and Fu, Y. (2022). Visual pigment-deficient cones survive and mediate visual signaling despite the lack of outer segments. Proc Natl Acad Sci U S A 119.

22. Demontis, G.C., Gargini, C., Paoli, T.G., and Cervetto, L. (2009). Selective Hcn1 channels inhibition by ivabradine in mouse rod photoreceptors. Invest Ophthalmol Vis Sci 50, 1948–1955.

23. Fain, G.L., Quandt, F.N., Bastian, B.L., and Gerschenfeld, H.M. (1978). Contribution of a caesium-sensitive conductance increase to the rod photoresponse. Nature 272, 466–469.

24. Ingram, N.T., Sampath, A.P., and Fain, G.L. (2020). Membrane conductances of mouse cone photoreceptors. Journal of General Physiology Mar 2;152(3):e201912520. doi: 10.1085/jgp.201912520.

25. Fain, G.L., Quandt, F.N., and Gerschenfeld, H.M. (1977). Calcium-dependent regenerative responses in rods. Nature 269, 707–710.

26. Fain, G.L., Gerschenfeld, H.M., and Quandt, F.N. (1980). Calcium spikes in toad rods. J Physiol (Lond) 303, 495–513.

27. Yu, W.Q., Grzywacz, N.M., Lee, E.J., and Field, G.D. (2017). Cell type-specific changes in retinal ganglion cell function induced by rod death and cone reorganization in rats. J Neurophysiol 118, 434–454.

28. Koch, K., McLean, J., Berry, M., Sterling, P., Balasubramanian, V., and Freed, M.A. (2004). Efficiency of information transmission by retinal ganglion cells. Curr Biol 14, 1523–1530.

29. Chichilnisky, E.J. (2001). A simple white noise analysis of neuronal light responses. Network 12, 199–213.

30. Torre, V., Straforini, M., Sesti, F., and Lamb, T.D. (1992). Different channel-gating properties of two classes of cyclic GMP-activated channel in vertebrate photoreceptors. Proc R Soc Lond B Biol Sci 250, 209–215.

31. Rieke, F., and Schwartz, E.A. (1994). A cGMP-gated current can control exocytosis at cone synapses. Neuron 13, 863–873.

32. Cha, S., Ahn, J., Jeong, Y., Lee, Y.H., Kim, H.K., Lee, D., Yoo, Y., and Goo, Y.S. (2022). Stage-Dependent Changes of Visual Function and Electrical Response of the Retina in the rd10 Mouse Model. Front Cell Neurosci 16, 926096.

33. Wang, T., Pahlberg, J., Cafaro, J., Frederiksen, R., Cooper, A.J., Sampath, A.P., Field, G.D., and Chen, J. (2019). Activation of Rod Input in a Model of Retinal Degeneration Reverses Retinal Remodeling and Induces Formation of Functional Synapses and Recovery of Visual Signaling in the Adult Retina. J Neurosci 39, 6798–6810.

34. Botto, C., Rucli, M., Tekinsoy, M.D., Pulman, J., Sahel, J.A., and Dalkara, D. (2022). Early and late stage gene therapy interventions for inherited retinal degenerations. Prog Retin Eye Res 86, 100975.

35. Jones, B.W., Pfeiffer, R.L., Ferrell, W.D., Watt, C.B., Marmor, M., and Marc, R.E. (2016). Retinal remodeling in human retinitis pigmentosa. Exp Eye Res 150, 149–165.

36. Yao, X., Cafaro, J., McLaughlin, A.J., Postma, F.R., Paul, D.L., Awatramani, G., and Field, G.D. (2018). Gap Junctions Contribute to Differential Light Adaptation across Direction-Selective Retinal Ganglion Cells. Neuron 100, 216–228 e216.

37. Field, G.D., Gauthier, J.L., Sher, A., Greschner, M., Machado, T.A., Jepson, L.H., Shlens, J., Gunning, D.E., Mathieson, K., Dabrowski, W., et al. (2010). Functional connectivity in the retina at the resolution of photoreceptors. Nature 467, 673–677.

38. Litke, A.M., Bezayiff, E.J., Chichilnisky, E.J., Cunningham, W., Dabrowski, W., Grillo, A.A., Grivich, M., Grybos, P., Hottowy, P., Kachiguine, S., et al. (2004). What Does the Eye Tell the Brain?: Development of a System for the Large-Scale Recording of Retinal Output Activity. IEEE Transactions on Nuclear Science 51, 1434–1440.

39. Field, G.D., Sher, A., Gauthier, J.L., Greschner, M., Shlens, J., Litke, A.M., and Chichilnisky, E.J. (2007). Spatial properties and functional organization of small bistratified ganglion cells in primate retina. J Neurosci 27, 13261–13272.

40. Shlens, J., Field, G.D., Gauthier, J.L., Grivich, M.I., Petrusca, D., Sher, A., Litke, A.M., and Chichilnisky, E.J. (2006). The structure of multi-neuron firing patterns in primate retina. J Neurosci 26, 8254–8266.

41. Betsch, B.Y., Einhauser, W., Kording, K.P., and Konig, P. (2004). The world from a cat’s perspective--statistics of natural videos. Biol Cybern 90, 41–50.

42. Strong, S.P., Koberle, R., de Ruyter van Steveninck, R., and Bialek, W. (1998). Entropy and Information in Neural Spike Trains. Physical Review Letters 80, 197–200.

43. Ma, S.-K. (1981). Calculation of entropy from data of motion. Journal of Statistical Physics 26, 221–240.

